# FoldBench: An All-atom Benchmark for Biomolecular Structure Prediction

**DOI:** 10.1101/2025.05.22.655600

**Authors:** Sheng Xu, Qiantai Feng, Lifeng Qiao, Hao Wu, Tao Shen, Yu Cheng, Shuangjia Zheng, Siqi Sun

## Abstract

Accurate prediction of biomolecular complex structures is fundamental for understanding biological processes and rational therapeutic design. Recent advances in deep learning methods, particularly all-atom structure prediction models, have significantly expanded their capabilities to include diverse biomolecular entities, such as proteins, nucleic acids, ligands, and ions. However, comprehensive benchmarks covering multiple interaction types and molecular diversity remain scarce, limiting fair and rigorous assessment of model performance and generalizability. To address this gap, we introduce FoldBench, an extensive benchmark dataset consisting of 1,522 biological assemblies categorized into nine distinct prediction tasks. Our evaluations reveal critical performance dependencies, showing that ligand docking accuracy notably diminishes as ligand similarity to the training set decreases, a pattern similarly observed in protein–protein interaction modeling. Furthermore, antibody-antigen predictions remain particularly challenging, with current methods exhibiting failure rates exceeding 50%. Among evaluated models, AlphaFold 3 consistently demonstrates superior accuracy across the majority of tasks. In summary, our results highlight significant advancements yet reveal persistent limitations within the field, providing crucial insights and benchmarks to inform future model development and refinement.

## Introduction

Accurate modeling of biomolecular complexes at atomic resolution is central to understanding molecular mechanisms and guiding rational drug discovery(1–4). Advances in deep learningbased structural prediction methods, such as AlphaFold 2 (5), AlphaFold-Multimer (6), ESMFold (7), and RoseTTAFold2 (8), have demonstrated remarkable success in predicting the structures of protein monomers and protein complexes. Building upon these achievements, AlphaFold 3 has recently been introduced to extend structural prediction capabilities beyond proteins alone, covering a broader range of biomolecules, including ligands, ions, nucleic acids, and chemically modified residues.

Despite these advancements, the training code and datasets of AlphaFold 3 are not publicly available, limiting community efforts to iterate on and extend the frameworks. In response, several opensource reproductions such as Boltz-1 (9), Protenix (10), Chai-1 (11) and HelixFold 3 (12) have emerged. However, their accuracy across different biomolecule classes has not been systematically compared because a cross-domain benchmark is still missing.

The difficulty is further increased by marked diversity among target types. Antibody–antigen interfaces, protein–ligand complexes, and nucleic acids are widely considered hard cases because of their conformational flexibility and the limited number of experimental structures available. Therefore, a broad evaluation that spans these diverse interactions is essential for mapping current capabilities and identifying priorities for future improvement.

Existing benchmarking studies (13–19) have generally addressed a single task, covered a limited range of methods and targets, and often overlooked how prediction accuracy changes with similarity between test cases and training data. Without unified dataset construction and evaluation criteria, published performances are hard to compare and may not reflect true generalisation to novel targets.

We therefore present FoldBench, a unified benchmark of 1,522 low-homology biological assemblies that covers six major interaction types, ranging from monomers to multiple classes of intermolecular interaction. FoldBench supports fair, head-to-head assessment of recent all-atom predictors and provides a transparent baseline for future methodological advances.

Our contributions can be summarized as follows:

- **Comprehensive dataset** We curate and release FoldBench, a low-homology benchmark that spans proteins, nucleic acids, ligands, and six major interaction types, enabling assessments that were previously impossible with task-specific datasets.
- **Systematic evaluation** We conduct an extensive, side-by-side comparison of five state-of-the-art all-atom predictors, AlphaFold 3 (AF3), Boltz-1, Chai-1, and HelixFold 3 (HF3), quantify how performance varies with training-set similarity and target class, and highlight strengths as well as persistent limitations.
- **Open resources** All benchmark data, evaluation code and reference results are freely available, offering the community an objective starting point for future method development and refinement.

## Results

### Overview of the benchmark

This benchmark aims to establish a comprehensive dataset to make fair comparisons of current all-atom structure prediction models. To prevent potential data leakage, we collected the first bioassemblies from PDB entries after 2023-01-13 (validation set cutoff of AlphaFold 3) and before 2024-11-01. Targets with high sequence or structural similarity to entries in the training set were removed (see Methods), resulting in a low-homology benchmark dataset. The curation process of the benchmark dataset is illustrated in Fig. 2A.

**Fig. 1.**
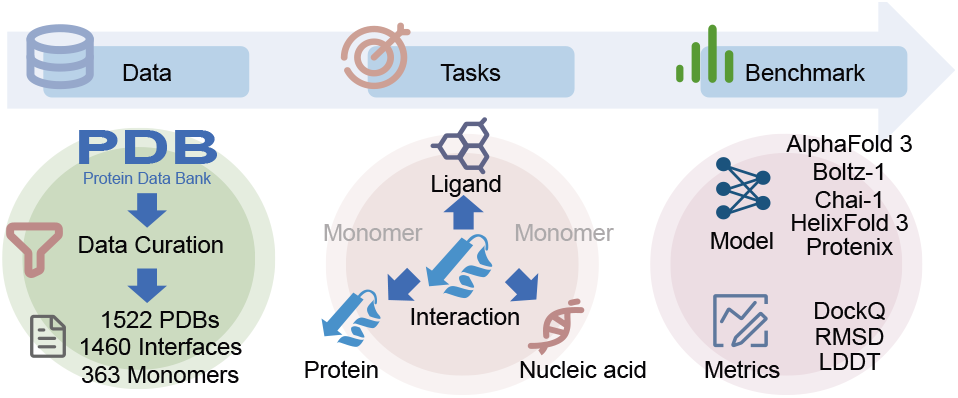
Overview of the benchmark workflow for evaluating all-atom structure prediction models.

**Fig. 2.**
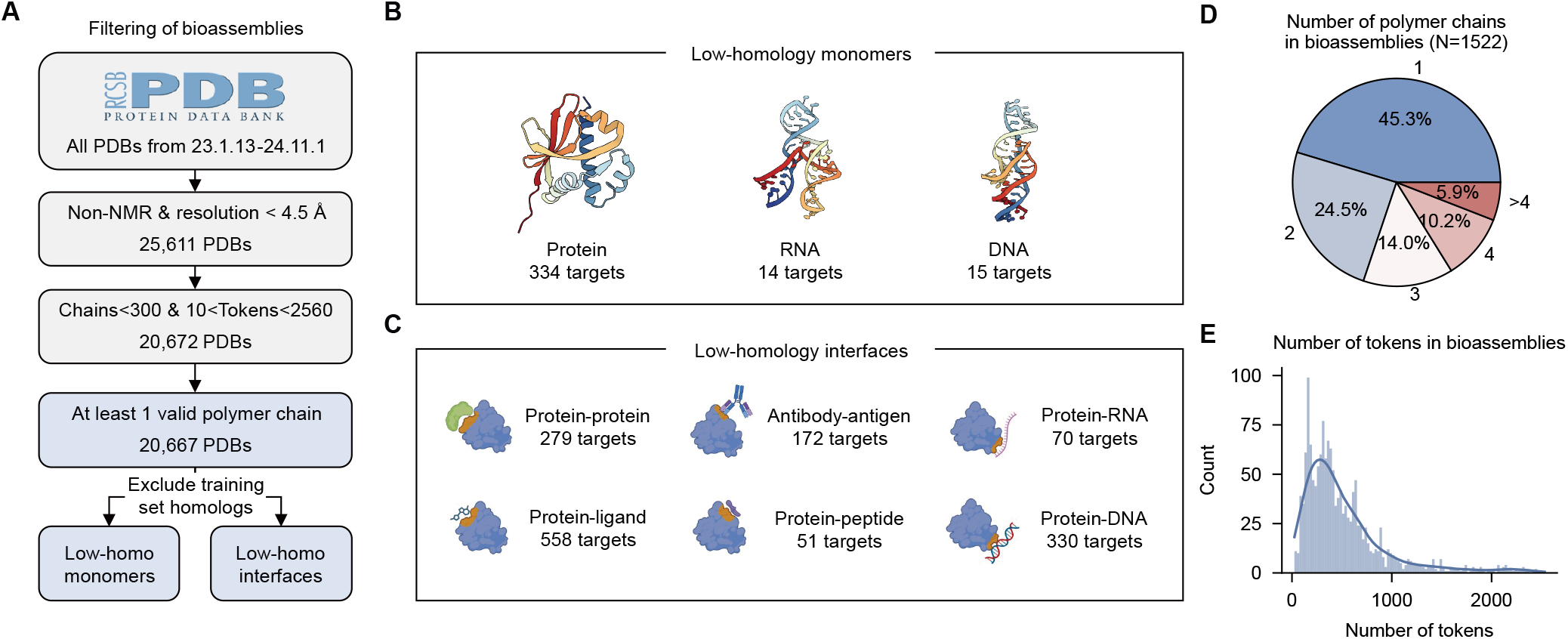
Data curation pipeline and statistical summary of the benchmark dataset. **A**. Workflow illustrating the selection and processing steps applied to biological assemblies retrieved from the Protein Data Bank (PDB). Resulting datasets comprise low-homology monomer and interface targets. **B**. Number of low-homology monomer targets categorized by biomolecular types. **C**. Number of low-homology interface targets categorized by interface types. **D**. Distribution of polymer chain counts across biological assemblies in the benchmark dataset (N=1,522). **E**. Distribution of token counts per biological assembly.

As shown in Fig. 2B-C, the benchmark dataset includes monomers and various interfaces, containing 334 protein monomers, 14 RNA monomers, and 15 DNA monomers, as well as 6 interface types involving 279 protein-protein, 172 antibody-antigen, 558 protein-ligand, 70 protein-RNA, 330 protein-DNA, and 51 protein-peptide interfaces. Fig. 2D-E provide the statistics of the dataset, the majority of complexes have two or fewer polymer chains, accounting for 69.8%, and most complexes have less than 1,000 tokens under AlphaFold 3’s tokenization scheme.

We used OpenStructure (20) as our primary evaluation tool. For most protein-related interfaces, we applied DockQ (21) to compute the docking accuracy. The DockQ success rate is the ratio of predictions achieving a DockQ score *≥* 0.23, the threshold indicating a successful docking pose (21).

Benchmark evaluation was performed on either individual monomer chains or distinct interfaces extracted from predicted full biological assemblies. Predictions were generated using a 5×5 sampling strategy (5 seeds × 5 samples) with 10 recycles to ensure thorough conformational space exploration (22) for each model. For more details, please refer to the Methods section.

### Benchmarking protein-ligand co-folding performance

Protein-ligand interactions are crucial for most pharmaceutical interventions (4, 23). Therefore, accurate prediction and characterization of these interactions are vital for modern drug discovery, facilitating the identification of small molecules that can specifically alter protein function (24). The emergence of AlphaFold 3 has been reported to be better than classical docking tools in specified tasks. (22, 25). For such co-folding methods, it is essential to assess the accuracy of both the ligand and protein structures (13). Fig. 3A illustrates the ligand success rate for five models, based on the overall dataset (558 targets), the scores for the “unseen protein” subset (76 targets), and the scores for the “unseen ligand” set (482 targets). Here, “unseen protein” refers to proteins in the protein-ligand pairs that exhibit less than 40% sequence identity to any protein sequence in the training set. Meanwhile, “unseen ligand” indicates that, although the protein sequence from the interface has homologs in the training set, the ligand must have less than 0.50 Tanimoto similarity to any ligands in complexes containing the homologous protein.

**Fig. 3.**
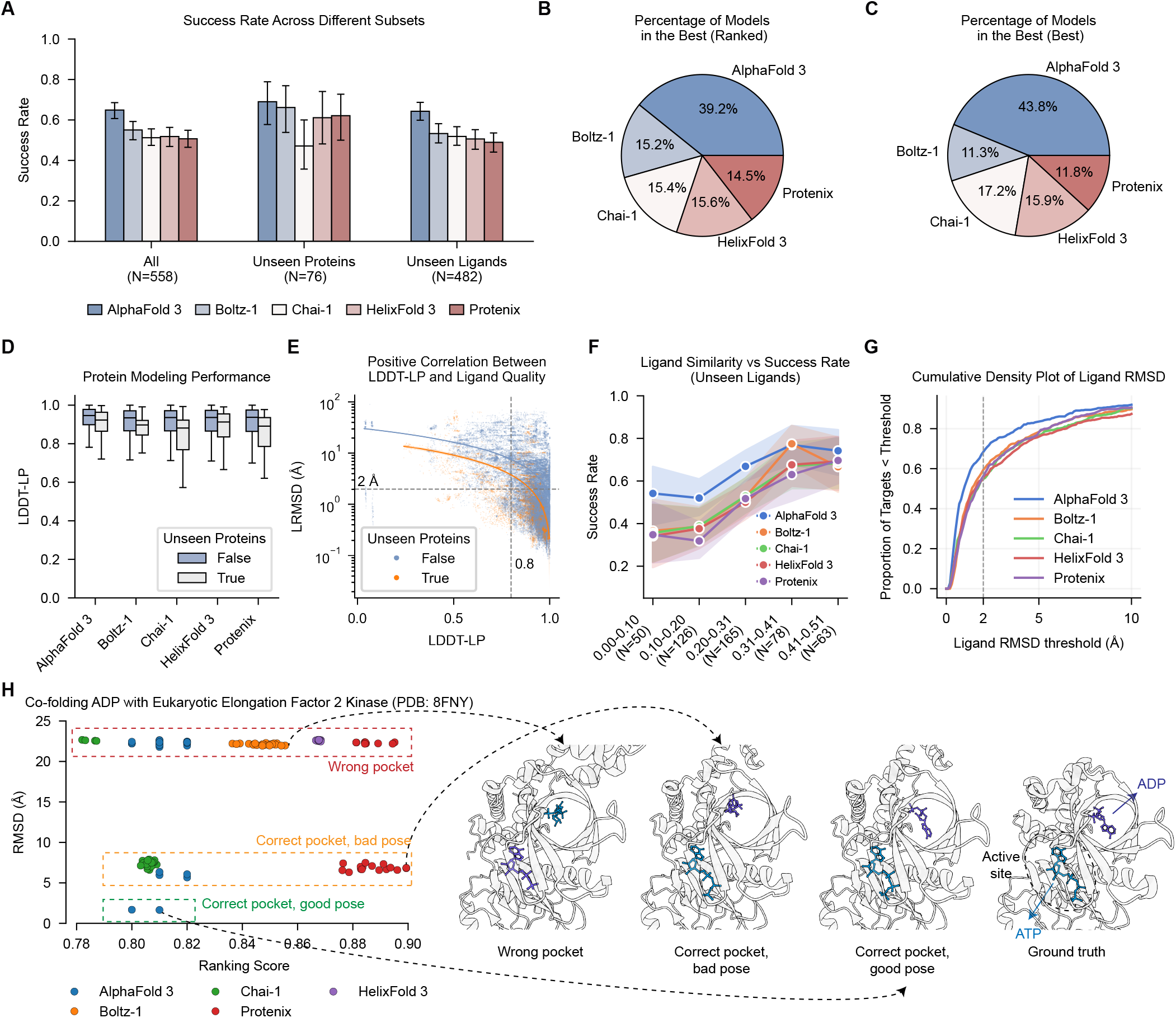
Performance on protein-ligand interaction modeling. The protein-ligand dataset can be divided into two categories: unseen proteins and unseen ligands. “unseen protein” means the protein in the protein-ligand pairs is unseen (e.g. less than 40% sequence identity to the training set), and “unseen ligand” means that we allow the protein sequence in the interface can have homologs in the training set, but the ligand should have less than 0.5 Tanimoto similarity to the ligands in the same complex where the protein homolog is in. **A**. Success rate across different subsets, where the success rate is defined as the ratio of LRMSD less than 2 Å and LDDT-PLI (Local Distance Difference Test for Protein-Ligand Interactions) > 0.8. **B**. The percentage of predictions made by each model is the best (lowest LRMSD among all models’ predictions). The predicted structure is selected by ranking score (left) and **C**. best scored (right). **D**. Protein (pocket) modeling quality for both low-homology and homologous. **E**. Scatter plot of protein pocket LDDT and LRMSD. **F**. Ligand similarity vs success rate in the unseen ligand subset. **G**. Cumulative Density of Ligand RMSD, varying from 0 to 10. **H**. Case study of co-folding ADP with eukaryotic elongation factor 2 kinase (PDB: 8FNY).

For the entire protein-ligand set, AlphaFold 3 achieves a 64.9% success rate, surpassing the runner-up, Boltz-1, by nearly 10%. Surprisingly, success rates rise across most predictors when we focus on the “unseen protein” subset, where AlphaFold 3 reaches 69.0%. By contrast, for the “unseen ligands” set, AlphaFold 3’s performance (64.3% success rate) is close to the performance on the overall dataset.

We first look into the protein modeling quality among the two sets to further investigate the performance gap between the unseen protein and the unseen ligand subset. Fig. 3D presents the lig- and pocket modeling performance for both low-homology and non-low-homology proteins, evaluated using LDDT-LP (ligand pocket LDDT). All methods perform well, with scores generally above 0.8, although low-homology proteins slightly impact the scores, which aligns with the good performance on protein monomers (Supplementary Note 1). We also showcase that protein modeling quality (LDDT-LP) has a positive correlation with ligand quality (Fig. 3E) for both the unseen protein subset (orange) and unseen ligand subset (blue). For targets having LDDT-LP greater than 0.8, their LRMSDs typically range below 2 Å for the unseen protein subset, as we did not impose a ligand similarity limitation on this subset. The same trend has been observed in the unseen ligand subset, but overall LRMSD is higher. It suggests that protein modeling difficulty is unlikely to be the primary bottleneck in protein–ligand interface modeling; instead, ligand similarity appears to be the more decisive limiting factor.

Fig. 3F examines the impact of ligand similarity to the training set on the modeling accuracies. The x-axis represents different ligand similarity ranges, while the y-axis shows the success rate of ligand docking. As similarity increases, the model’s ability to accurately model ligands also improves, with AlphaFold 3 showing a slight performance advantage. This trend indicates that models rely substantially on memorized ligand binding modes encountered during training. In practice, this means that current all-atom predictors are effective at recapitulating known protein–ligand complexes but exhibit limited generalization to unseen ligands.

We analyzed the proportion of targets with RMSD values below different thresholds, as shown in Fig. 3G, with the traditional ligand RMSD threshold of 2.0 indicated by a dashed line. It is clear that AlphaFold 3 consistently outperforms other models across various thresholds, while the performance of the other models shows no significant distinction.

Fig. 3B–C report, for each method, the fraction of targets whose lowest ligand-RMSD (LRMSD) among the 25 predictions attains the overall best value. In panel B, predictions are ordered by each model’s ranking score; in panel C, they are ordered by the LRMSD against the ground truth. Under both ranking schemes, AlphaFold 3 takes the lead, providing the best prediction for roughly 40% of the targets.

Allosteric regulation occurs when a ligand binds to a site topographically distinct from the orthosteric active site, inducing conformational or dynamic changes that modulate protein function without directly competing with endogenous substrates (26). This modulation mode often affords greater receptor subtype selectivity and reduced liability for off-target effects. Computational identification of allosteric sites is therefore critical for structure-based drug design.

As shown in Fig. 3H, PDB entry 8FNY reveals ADP bound within a distinct pocket situated on the face opposite to the kinase’s orthosteric ATP-binding site (27). In this case, most prediction results were incorrect, including both inaccurate pocket predictions and correct pocket predictions with incorrect conformations. Only two predictions from AlphaFold 3 were accurate. However, while there are successful predictions among AlphaFold 3’s 25 samples, it is hard to distinguish them by the ranking scores of the model. This ranking bias likely reflects the overrepresentation of orthosteric complexes in the training data and the high chemical similarity between ATP and ADP. Moreover, the steric constraints of this distal site—whose geometry is incompatible with an ATP (27)—underscore the necessity of incorporating non-orthosteric binding examples and refining ranking algorithms to improve recognition of allosteric interactions.

Since 8FNY has a small molecule at the orthosteric site, we aim to further investigate the model’s performance on allosteric sites when no ligand is present at the orthosteric site. Consequently, we selected the popular target CDK2 (PDB: 7RWF (28)) for our study. As illustrated in Fig. S3, although the protein structures are correctly predicted, the ligand docking quality remains incorrect, with LRMSD mostly exceeding 10 Å. All models incorrectly placed the ligand in the active site where ATP should be positioned.

### Modeling protein-protein interactions

Protein-protein interactions (PPIs) mediate almost every cellular process, from signal transduction to metabolic regulation. Structurally characterizing these interfaces is essential for understanding biological mechanisms and developing therapeutic PPI modulators (29, 30).

Fig. 4A illustrates the relationship between complex LDDT and structural similarity against the training set for the initial 501 targets, before structure redundancy filtering. Prediction accuracy rises steadily as structural similarity increases, suggesting a potential structural-memory effect in the models.

**Fig. 4.**
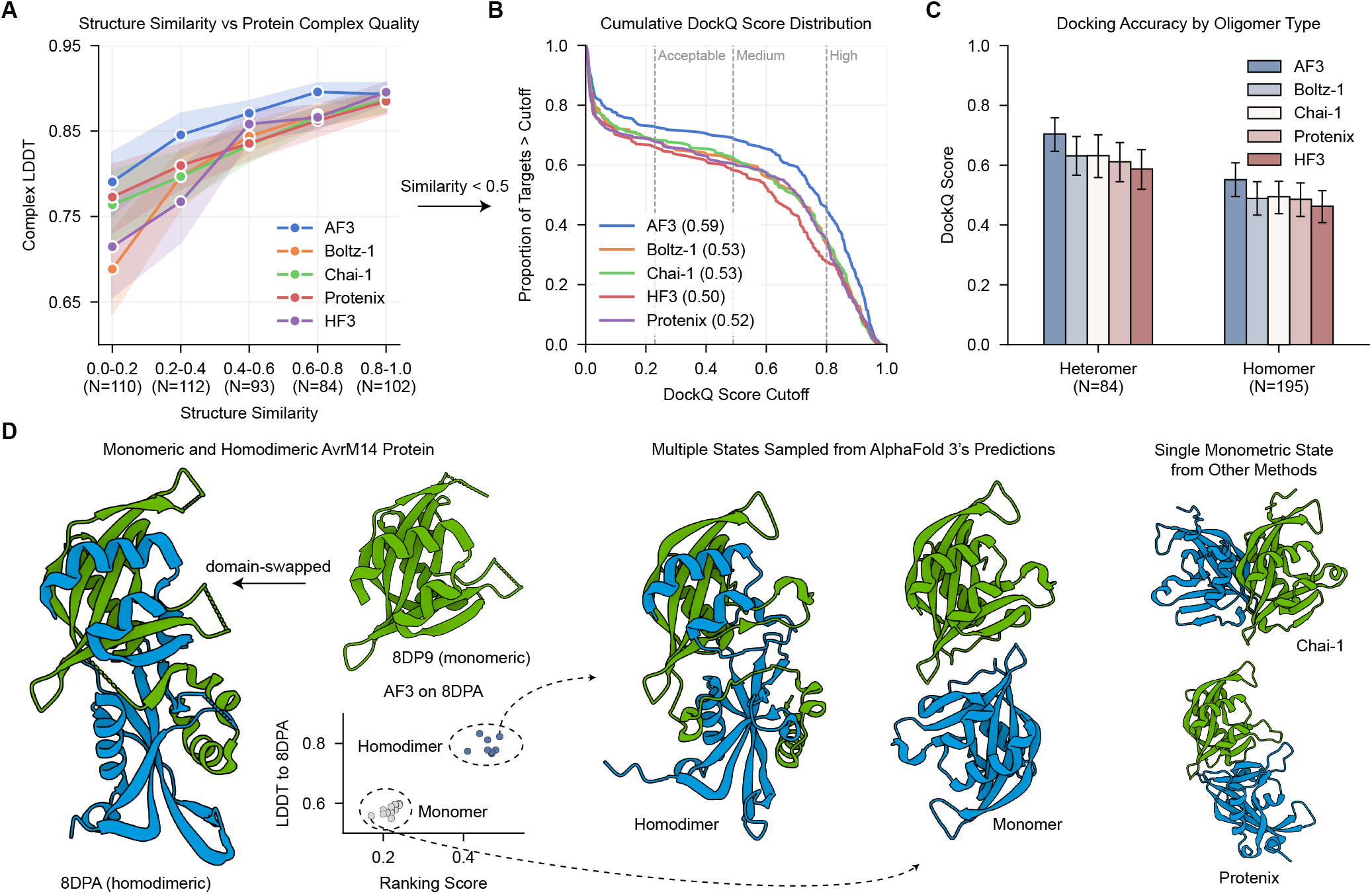
Model performance on protein-protein (N=279, without antibody-antigen) task. **A**. Complex LDDT varies with structural similarity against the training set. **B**. Cumulative distribution of DockQ scores for the same 279 targets. **C**. Model performance separated by oligomeric state: heteromer (N=84) and homomer (N=195). **D**. Case study: the AvrM14-B Nudix hydrolase effector (PDB 8DPA, homodimeric). AlphaFold 3 sampling produced both the experimentally observed domain-swapped homodimer and a monomeric state, and ranking scores can clearly separate the two basins (scatter plot). Competing methods (right) converge exclusively on the monomeric state.

To alleviate potential model memorization and to incorporate more challenging targets, we therefore constituted our proteinprotein target subset exclusively from targets exhibiting low structural similarity, defined as a TM-score < 0.5 relative to complexes in the training set. After filtering out these targets, our final proteinprotein subset comprises 279 protein-protein interfaces.

Figure 4B presents the cumulative DockQ-score distributions for these 279 targets as the success rate threshold ranges from 0 to 1. AlphaFold 3 consistently leads all other models, achieving an AUC of 0.63 compared to 0.57 (Boltz-1), 0.56 (Chai-1), 0.55 (Protenix) and 0.54 (HelixFold 3). At the conventional success threshold of 0.23, AlphaFold 3 attains a 73.7% DockQ success rate—over 5 percentage points higher than the runner-up, Chai (68.5%) (Table S3).

After splitting by oligomeric state, heteromeric interfaces show higher mean DockQ scores than homomeric ones (Fig. 4C), opposite to the trend reported for AlphaFold-Multimer (6). A likely reason is that, in heteromers, high global structural similarity to the training set does not necessarily preserve the correct subunit orientation, so docking can fail even when the complex TM-score is high (Fig. S5).

Figure 4G highlights AlphaFold 3’s potential to model conformational changes in complexes. For the domain-swapped homodimer 8DPA (31), it not only predicts the swapped conformation accurately but also ranks the correct dimer highest among 25 candidates. However, the other models fail to reproduce the dimeric conformation and even misidentify the docking interface. Future work should focus on developing learning strategies that extract maximal information from the limited number of experimentally determined structures exhibiting conformational changes, thereby improving the models’ ability to generalize to flexible structural rearrangements.

### Results on antibody-antigen complexes

Antibody-antigen interactions form the molecular foundation of adaptive immunity and represent critical targets for structural prediction due to their therapeutic implications (32, 33). Structural characterization of these complexes provides essential insights for epitope mapping, rational antibody engineering, and accelerated therapeutic development. Despite computational advances, accurate prediction remains challenging due to CDR (complementarity-determining region) variability, binding-induced conformational changes, and diverse recognition modes (34, 35). These interfaces constitute particularly difficult targets in our benchmark, highlighting a frontier where improved modeling could significantly impact drug discovery timelines and success rates.

To analyse the model performance on antibody-antigen tasks, we benchmarked the models on 172 antibody-antigen pairs, including 123 antibodies, 46 nanobodies and 3 single-chain variable fragments (scFvs). Fig. 4A shows the cumulative distribution of DockQ scores for each model. Compared to the performance on proteinprotein interface, the success rate on antibody-antigen dropped to 47.9% for AlphaFold 3, while other methods exhibited over 60% of failure rate. Despite the modest overall performance, AlphaFold 3 still takes the lead at most DockQ score cutoffs (AUC=0.36), significantly outperforming the second-best model (Protenix) by 0.13.

Given that AlphaFold 3 achieves enhanced performance for antibodies through extensive sampling (1,000 seeds) (22), we aimed to assess the impact of a deeper sampling space. As shown in Fig. 5B, we progressively increased the number of samples in the order of seeds, selecting the top-ranked sample by ranking score at each increment for evaluation. The success rate of AlphaFold 3 gradually increases with more samples, demonstrating the advantages of increased sampling. However, other models exhibit greater fluctuations and even declines, indicating that increasing the number of samples without robust ranking capabilities may result in ineffective sampling and lower-quality conformations. To fully leverage the benefits of increased sampling and thereby obtain the best possible sampled conformations, the ranking methodology is required to approximate “oracle” (i.e., identifying the best-scored prediction) performance, as depicted in Fig. S5A.

**Fig. 5.**
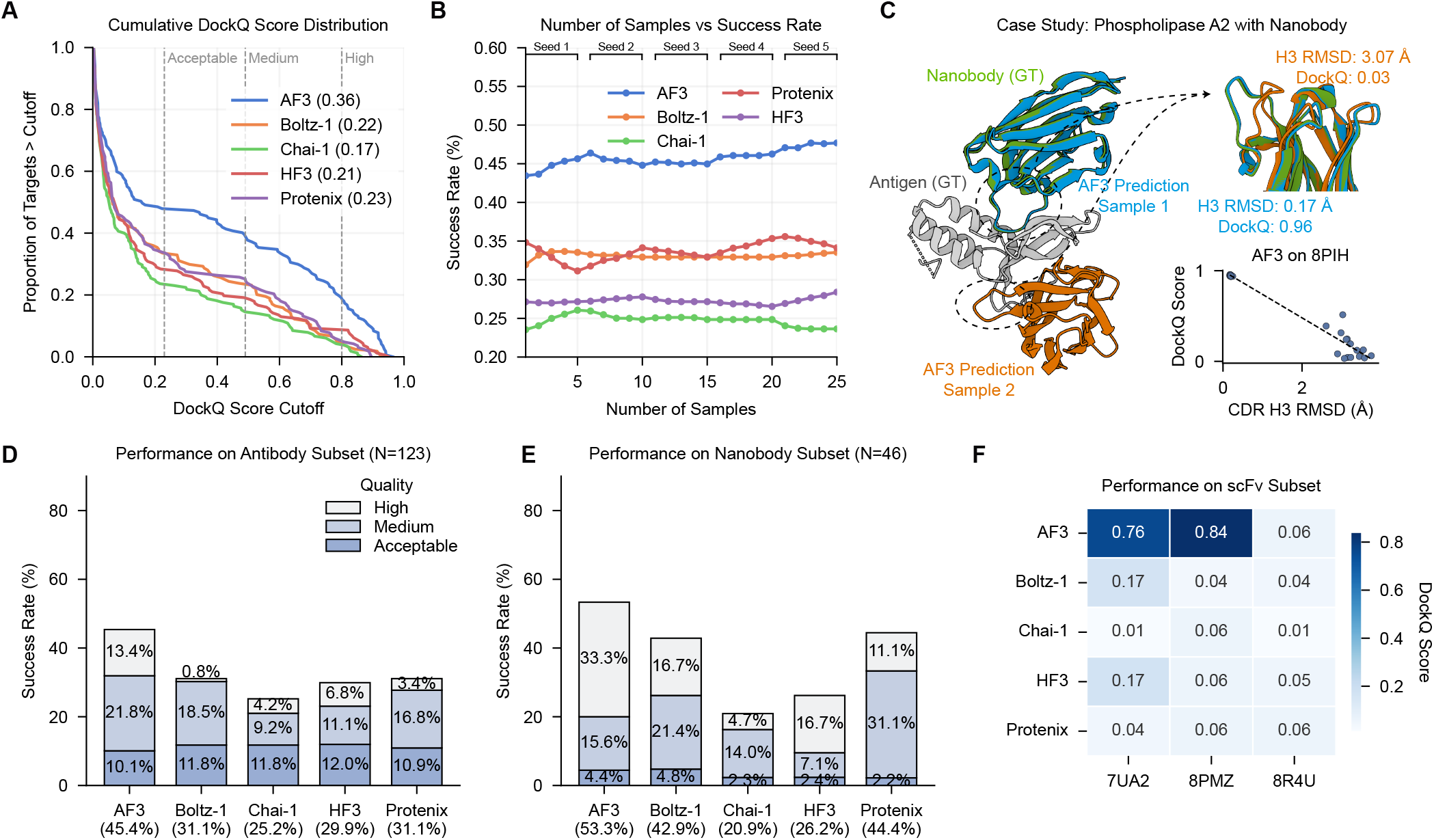
Model performance on antibody-antigen (N=172) subset. **A**. Cumulative distribution of DockQ scores for the 172 targets. **B**. DockQ average success rate by number of samples. **C**. Case study: The relationship between CDR H3 loop modeling quality and docking performance. **D**. Docking performance on standard antibody-antigen subset (N=123). **E**. Docking performance on nanobody-antigen subset (N=46). **F**. DockQ score on scFv subset (N=3).

Additionally, we explored the effect of increasing the number of generated samples through two primary methods. The first approach involved using different random seeds, where each seed initiated a complete, independent run of the entire pipeline, encompassing both backbone generation and the diffusion process. In contrast, the second approach consisted of a single execution of the initial backbone extractor, with its output then serving as the condition for multiple diffusion samples. The results of the two strategies are shown in Fig. S5B. When a ranking score was employed to identify potential successes, the two strategies did not exhibit a significant difference in performance. However, if an “oracle” selection was applied, the former strategy (full re-execution with diverse seeds) yielded markedly superior results. This suggests that while the full re-execution approach is more computationally expensive, it facilitates a broader exploration of diverse, high-quality structure predictions; yet, these better predictions are not guaranteed to be selected by the model’s ranking procedure.

The case study in Fig. 5C further illustrates the critical influence of CDR H3 loop modeling quality on antibody-antigen docking accuracy. We select the nanobody-phospholipase A2 complex (PDB: 8PIH) as an example, with the experimental structures of the antigen and nanobody shown in grey and green, respectively. Two predictions from AlphaFold 3 are highlighted: one (Sample 1, blue) exhibits near-native H3 conformation with an H3 RMSD of 0.17 Åand achieves a high DockQ score of 0.96, indicating a highly accurate binding. In contrast, another sample (Sample 2, orange) shows an incorrect H3 conformation (H3 RMSD: 3.07 Å), resulting in a poor DockQ score of 0.03. The scatter plot further quantifies this relationship: for 25 samples, DockQ scores decrease as the H3 RMSD increases. This strong anti-correlation underscores the role of precise CDR H3 loop modeling in contributing to overall docking success.

To further analyze the performance of different antibody types, the antibody-antigen interactions are split into three groups: standard antibodies (N=123), nanobodies (N=46) and scFvs (N=3). Fig. 5D shows the overall model docking performance on antibodies, measured by the DockQ success rate. Most models performed poorly, with only AlphaFold 3 exceeding a 40% success rate at 45.4%. Other models averaged around 30%, highlighting the challenges of the antibody-antigen prediction task. In terms of highquality docking, AlphaFold 3 also excelled, achieving a score of 13.4%, indicating its ability to make accurate predictions while ensuring high quality. In contrast, the proportion of high-quality docking in other models is significantly lower, with Boltz-1 scoring just 0.8% for high-quality docking.

Relative to conventional antibodies, nanobodies present a more compact, single-domain binding interface. In this setting, AlphaFold 3, Boltz-1 and Protenix achieve substantially higher docking success (Fig. 5E): AlphaFold 3 reaches a 53.3 % overall success rate with 33.3% of predictions classified as high quality. By contrast, Chai-1 and HelixFold 3 attain only 20.9% and 26.9%, respectively. This performance gap likely reflects the advantage of reduced inter-chain complexity for methods with accurate loop modeling and ranking routines.

Single-chain variable fragment (scFv) is a type of engineered antibody that consists of the variable regions of the heavy and light chains of an antibody connected by a short linker. Fig. 5F presents the DockQ score average performance on scFv in our benchmark, including 7UA2, 8PMZ, and 8R4U. In this case, AlphaFold 3 outperformed the others, successfully predicting 2 out of 3 instances, while the other models failed to make any correct predictions. This suggests that the limited training data for scFv may limit the performance of the models.

### Assessment of nucleic acids

Resolving structures of nucleic acids is crucial for understanding biological processes (36). A significant advancement made by these all-atom structure prediction models is their capability to model nucleic acids, whereas earlier approaches often required developing dedicated architectures for RNA or DNA tasks (37–39). Our results demonstrate that nucleic acid structure prediction remains a major challenge for all current models. As shown in Fig. 6A, C, the average LDDT scores for RNA and DNA monomers typically fall within the range of 0.2 to 0.6, markedly lower than those for protein monomers (up to 0.88). For both tasks, AlphaFold 3 achieves the highest performance (LDDT 0.53 for DNA and 0.61 for RNA) among the five models, followed by Proteinx (0.44 for DNA and 0.59 for RNA) and Chai-1 (0.46 for DNA and 0.49 for RNA). Although HelixFold 3 has a relatively high LDDT in RNA monomers (0.55), its accuracy on DNA is reduced (0.29), primarily due to the poor performance of G-quadruplexes (e.g., PDB: 8D78 and 8P6B). Boltz-1 exhibits similar performance on these structures as well.

**Fig. 6.**
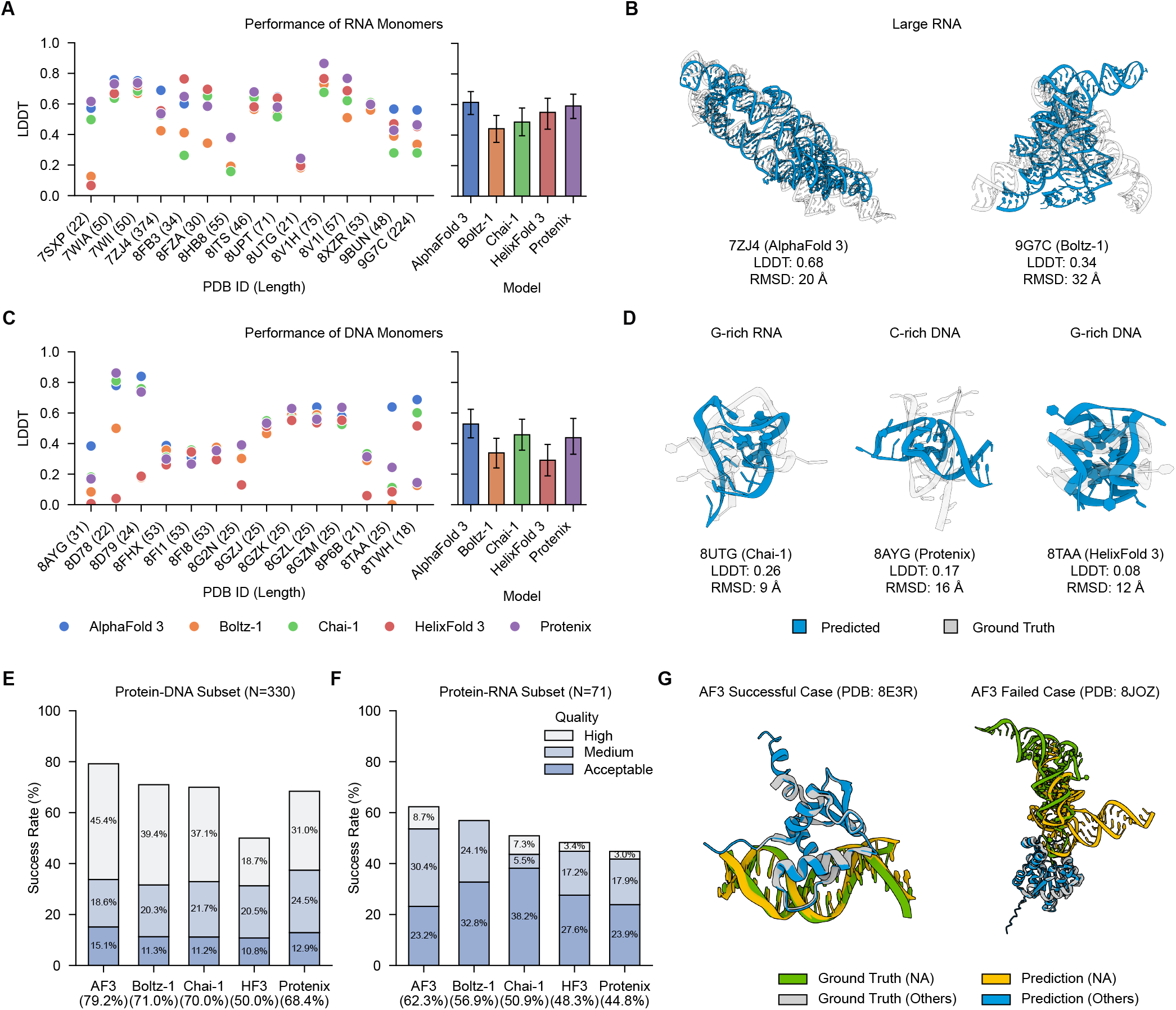
Summary of performance of different models on nucleic acids prediction tasks. **A**. Performance on RNA monomer tasks, detailed comparison of each target and summary respectively. **B**. Visualization of large RNA targets (grey) and predicted structures (blue), where most of the models failed. **C**. Performance on DNA monomer tasks, detailed comparison of each target and summary, respectively. **D**. Performance on RNA monomer tasks, detailed comparison of each target and summary respectively. **D**. Visualization of G/C-rich nucleic acid targets (grey) and predictions (blue), which are also acknowledged as hard targets. **E**. Docking performance on protein-DNA subset (N=330). **F**. Docking performance on protein-RNA subset (N=71). **G**. Visualization of successful (Protein-dsDNA) and unsuccessful (Protein-tRNA) cases of AlphaFold 3.

The prediction of large or highly flexible nucleic acids poses particular difficulties. For instance, in the case of an RNA aptamer (PDB: 7ZJ4, length=374) and a non-coding RNA (PDB: 9G7C, length=224), all models fail to reconstruct the global architecture (Fig. 6B). This is evidenced by high LDDT scores (e.g., 0.68 for 7ZJ4 by AlphaFold 3) alongside extremely large RMSD values (20 Å). This discrepancy indicates that while local elements like hairpins or regular helices can be accurately captured, the modeling of tertiary folds and long-range interactions remains largely unsolved.

Another challenging target type is G/C-rich nucleic acids (Fig. 6D). For example, PDB entry 8UTG is an RNA from the NS5 gene in the West Nile Virus genome, with a short length of 21 nucleotides. Although 8UTG is not very long, its G-Quadruplex structure still poses certain challenges for the model. The visualized cases illustrate that the model struggles to learn G/C-rich complexes, frequently with low LDDT and high RMSD.

Fig. 6E-F shows the performance on protein-DNA and protein-RNA interactions. AlphaFold 3 continues to perform the best, achieving a success rate of 79.2% in the protein-DNA task. However, the scores for protein-RNA interactions are lower than those for protein-DNA, with AlphaFold 3’s score dropping to 62.3%, and all models showing a significant decrease in the proportion of highquality predictions. Fig. 6G illustrates both successful (Protein-dsDNA) and unsuccessful (Protein-tRNA) cases of AlphaFold 3. Compared to the more structured system in protein-dsDNA complexes, the inherent complexity and flexibility of nucleic acid structures might make predictions more challenging for these structure prediction models.

The underperformance of current models on nucleic acid structure prediction primarily stems from data scarcity. As of May 2025, nucleic acid structures account for only about 8% of the 237,000 entries in the PDB, severely limiting the diversity and generalizability that models can achieve compared to protein-rich datasets. Beyond data constraints, nucleic acids—particularly RNA—display remarkable structural heterogeneity and conformational flexibility, including non-canonical base pairs, complex tertiary motifs, and dynamic long-range interactions, all of which compound the intrinsic modeling difficulty. Furthermore, nucleic acid folding and stability depend highly on environmental factors such as ion concentration and pH (40), variables that most current prediction algorithms do not explicitly encode. The relatively higher performance on protein-DNA complexes can be attributed to the greater structural regularity of DNA (41), especially in canonical double helices, as well as to the richer training data available for DNA-binding proteins. By contrast, protein-RNA interactions remain particularly challenging due to the pronounced structural diversity (42) and limited structural data available for RNA.

## Conclusions

To comprehensively evaluate existing models capable of general all-atom structure prediction across various biomolecular systems, we established FoldBench. This systematic benchmark comprises 1,522 biological assemblies spanning 9 target types. The assessment of AlphaFold 3, Boltz-1, Chai-1, HelixFold 3, and Protenix across these diverse systems reveals significant advancements alongside persistent challenges within the field of structural prediction.

AlphaFold 3 consistently outperforms other models across all evaluation metrics and structural categories. Its superior abilities in monomer and interaction prediction, conformational change modeling, and ranking underscore its remarkable generalization and robustness, positioning it as the leading model.

However, inherent weaknesses persist across these models. All models exhibit a degree of memorization, an issue common in data-driven approaches. While most models perform well on tasks with sufficient data or relatively simple structures (e.g., protein monomers), their performance declines in data-scarce domains. This is particularly apparent for nucleic acids and for predicting specific sites or conformers crucial in drug discovery, such as allosteric sites in protein-ligand interactions and cyclic peptides (Supplementary Note 2).

Compared with protein–protein interfaces, antibody–antigen interactions remain challenging. Accurate modeling of CDRs is particularly difficult owing to their intrinsic conformational flexibility and the absence of homologous sequence templates. Since CDR loops lack conserved homologues, MSA-based methods provide little guidance for their conformations, and errors in CDR modeling critically impair docking success. Moreover, our antibody–antigen analysis shows that even extensive sampling yields limited benefit when ranking functions cannot reliably select near-native CDR conformations.

These findings also suggest potential future research directions. For the research community, expanding the scale and diversity of foundational training datasets, coupled with more active data sharing, is essential. Furthermore, exploring data synthesis techniques, such as self-distillation, could offer valuable strategies to augment available data. Regarding ranking, the development of more effective scoring functions or ranking algorithms appears crucial, with potential exploration of contemporary approaches such as reinforcement learning.

## Methods

### Data curation procedure

This benchmark aims to create a low-homology and comprehensive dataset for fair comparison of current all-atom bimolecular structure prediction models. To remove homology targets in the benchmarks set, we first reproduced the training set of AlphaFold 3 (2021-09-30 cutoff), and constructed the benchmark dataset as follows (see also Fig. 2A).

Filtering of targets:

- All entries must be released on PDB (Protein Data Bank) after 2023-01-13 (AlphaFold 3 validation cutoff), and before 2024-11-01.
- Entry must be non-NMR and with a resolution of less than 4.5 Å.
- Number of polymer chains should be less than 300.
- Number of tokens should be less than 2560 and larger than 10 under AlpaFold 3’s tokenization scheme.
- The first bioassembly of each entry was selected as the prediction target.

Filtering of bioassemblies:

- Water is removed.
- Polymer chains with less than 4 resolved residues are removed.
- Polymer chains with all unknown residues are filtered out.
- Protein chains with continuous *C*^*α*^ distance greater than 10Å are removed.
- Clashing chains are filtered out. For any pair of chains sharing >30% of all non-hydrogen atoms closer than 1.7 Å, the chain with the higher clash fraction (or, if equal, the smaller atom count) is removed.

Filtering to the low-homology set. We process targets into two types, monomer and interface (pairs of chains with minimum heavy atom (i.e., non-hydrogen) separation less than 5Å):

- **Monomers**: Monomer targets are bioassemblies that contain a single polymer (protein/DNA/RNA) chain, and have less than 40% sequence identity to the training set.
- **Polymer-polymer interface**: If both polymers have greater than 40% sequence identity to two chains in the same complex in the training set, then this interface is filtered out. For protein-protein interfaces, we further applied Foldseek-Multimer (43) to filter the interfaces, using a threshold of TM-score < 0.5 with the training set to exclude structurally similar ones.
- **Protein-ligand interface**: If the protein has greater than 40% sequence identity with a chain in a training complex and the ligand has greater than 50% Tanimoto similarity to a ligand from the sample complex, then the interface is filtered out. For further filtering, we randomly selected a single interface as the target for each ligand.
- **Protein-peptide interface**: For interfaces between a protein and a peptide (less than 16 residues), the protein chain should have less than 40% sequence identity to the training set.

Final clustering and resolution cut-offs. The remaining low-homology assemblies were clustered (MMseqs2 for proteins at 40 % identity; 100 % identity for nucleic acids and peptides). Within each cluster, the structure of the highest resolution was retained (excluding protein-ligand interfaces), subject to task-specific resolution limits:

- **Protein monomers**: 334 targets (7 de novo designed proteins were removed).
- **DNA monomers**: 15 targets.
- **RNA monomers**: 14 targets.
- **Protein–ligand interfaces**: 558 targets after additional ligand-quality filters (Supplementary Note 3).
- **Antibody–antigen interfaces**: 172 targets; resolution *<* 2.5 Å.
- **General protein–protein interfaces**: 279 targets (antibody-antigen complexes excluded); resolution *<* 2.0 Å.
- **Protein–peptide interfaces**: 51 targets; no further resolution constraint owing to limited data.
- **Protein–RNA interfaces**: 70 targets; resolution *<* 2.5 Å.
- **Protein–DNA interfaces**: 330 targets; resolution *<* 2.5 Å. Unless noted otherwise, performance was assessed on individual monomer chains or on the specified interfaces extracted from full-assembly predictions.

### Model Inference

First, we generated the JSON inputs of AlphaFold 3 by parsing the mmCIF files of the bioassemblies using the *folding_input*.*Input*.*from_mmcif* function from the AlphaFold 3 repository (https://github.com/google-deepmind/alphafold3) and removed bond information. Next, these JSON inputs were converted to other formats compatible with the different models, according to their respective input requirements and instructions.

After input generation, each model performed predictions using a 5×5 sampling strategy (5 seeds × 5 samples) with 10 recycles to ensure thorough conformational space exploration. Unless otherwise specified, the results are based on the structure with the highest ranking score for each model. Table S2 shows the commit ID and MSA source of each method. Inference was made on NVIDIA H800 80GB GPUs.

To predict more targets in the benchmark as much as possible, some modifications were made to run each prediction model:

For AlphaFold 3, input files with sequences that have no template hits will trigger a *StopIteration* error. We fixed this problem according to the related issue #364 (https://github.com/google-deepmind/alphafold3/issues/364). Note that AlphaFold 3 and HelixFold 3 include RNA MSA searching and template searching in their local searching pipeline, which differs slightly from the other three methods.

For Boltz-1, glycans, which were formatted as multiple Chemical Component Dictionary (CCD) entries corresponding to a single chain ID—were removed from the bioassemblies, as Boltz-1 cannot process this input structure.

Note that in our assessment version of the Chai-1 model, it cannot infer biological assemblies with a token count exceeding 2048, which will hinder the evaluation of those targets.

For HelixFold 3, there are nested *ProcessPoolExecutors* in the MSA searching pipeline, which easily get stuck in our practice, potentially due to memory limitations. We turn *ProcessPoolExecutors* into *ThreadPoolExecutors* limited by max_workers, which can help alleviate this issue. Additionally, since HelixFold 3 cannot handle modifications, we removed them from the input.

For Protenix, Protenix output CIF (Crystallographic Information File) doesn’t have _entity.type information, which OpenStructure uses to distinguish between polymer and non-polymer entities in the file, and we add this value extracted from the CIF information.

### Evaluation and Metrics

We used the widely adopted OpenStructure v2.8 as the main assessment tool to calculate the scores between the predicted results and the ground truth. For common monomer metrics such as LDDT, TM-score, and GDT-TS, we also use OpenStructure for calculations.

For most of the protein interface measurements, we used DockQ to compare the prediction with the ground truth complexes. The DockQ success rate is the ratio of predictions achieving a DockQ score greater than or equal to 0.23, the threshold indicating a successful docking pose (21). This score reflects the quality of the predicted interaction, with higher values suggesting a more reliable model. To provide a clearer assessment of docking results, DockQ scores can be divided into 4 bins (44):

- **Incorrect**: DockQ < 0.23
- **Acceptable**: 0.23 <= DockQ < 0.49
- **Medium**: 0.49 <= DockQ < 0.80
- **High**: DockQ > 0.80

The DockQ score for each interface was calculated using Open-Structure’s *compare-structures*, selecting based on the two native label chain IDs. However, OpenStructure does not support the DockQ score of the protein-nucleic interface, so we used the DockQ v2 program (44) as our assessment tool instead in this case. After completing the above adjustments and processing, Table S2 shows the final counts of predictable and assessable targets across various tasks and models, and the detailed performance results of various tasks and models are shown in Table S3.

The protein-ligand interface is measured using Binding-Site Superposed, Symmetry-Corrected Pose Root Mean Square Deviation, referred to as LRMSD (ligand RMSD) (45). Following (13), the lig- and docking success rate is defined as LRMSD < 2 Å and LDDT-PLI > 0.8. The LRMSD and LDDT-PLI scores for each proteinligand interface were calculated using OpenStructure’s *compareligand-structures*, selecting based on the two native label chain IDs. For antibody CDR loop assessment, we followed the evaluation pipeline outlined in (34) to calculate the CDR H3 loop RMSD. We renumbered the antibody structures using the AbNum web-server (46) according to the Chothia scheme (47), and then computed the H3 RMSD score using PyRosetta4 (48).

## CODE AVAILABILITY

All benchmark data, evaluation code and reference results can be accessed at https://github.com/BEAM-Labs/FoldBench.

## Supplementary Information

### Supplementary Note 1: Protein monomer performance

**Fig. S1.**
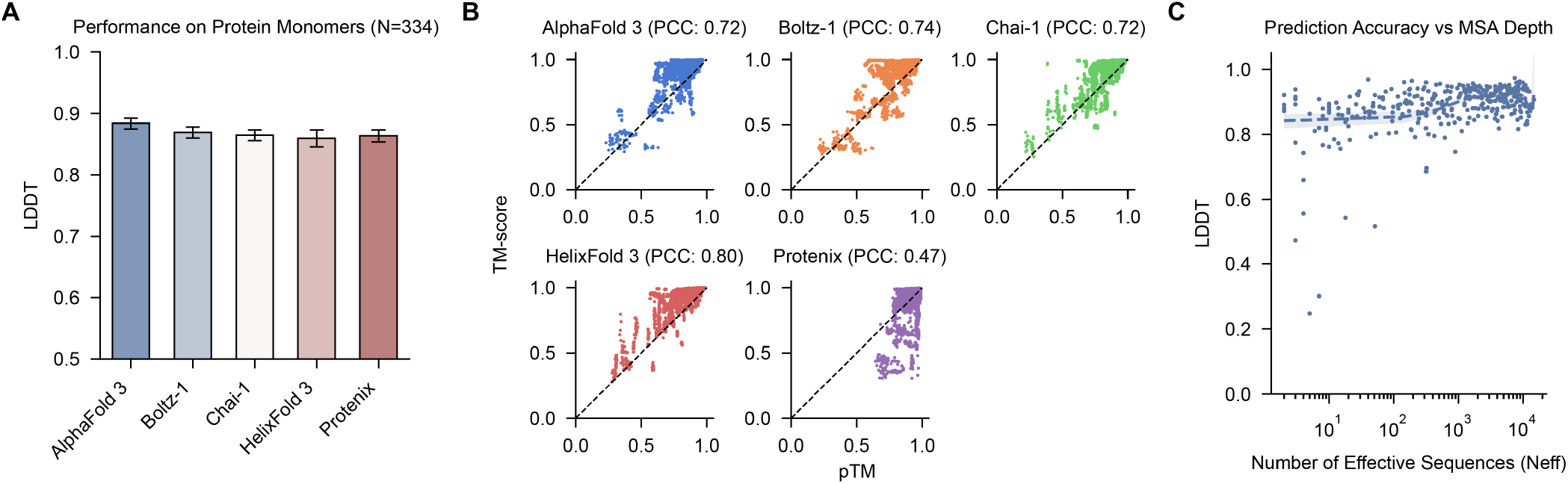
Summary of performance of different models on protein monomer prediction tasks. **A**. Performance comparison on protein monomers across different methods, with a sample size of 334 protein monomers. **B**. Pearson correlation coefficients between PTM and TM-Score across different methods in the protein monomer prediction task. **C**. The relationship between MSA depth (number of effective sequences, *N*^*eff*^ of the protein targets and LDDT.

Accurate prediction of monomeric structures forms the foundation for all interaction modeling. To evaluate protein structure prediction accuracy, we tested each method on the protein monomer dataset (N=334, excluding de novo designed proteins). Fig. S1A summarizes prediction quality in terms of local distance difference test (LDDT). The mean LDDT across targets exceeds 0.85 for every model, with AlphaFold 3 leading at 0.88, and no significant differences observed among the rest of the models, indicating all five models achieve a high protein monomer modeling quality.

High structural accuracy should be paired with well-calibrated error estimates if a predictor is to be used autonomously. Fig. S1B compares the predicted TM-score (pTM) with the observed TM-score for the 25 samples (5 seed × 5 diffusion samples) produced per target. AlphaFold 3, Boltz-1, Chai-1, and HelixFold 3 show Pearson correlation coefficients (PCC) between 0.72 and 0.80, indicating that their self-assessment captures true modeling quality in most cases. By contrast, Protenix produces over-confident scores (PCC=0.47), models with TM-score < 0.7 are frequently assigned pTM > 0.9. This mismatch would misrank candidates in large-scale screening or iterative design processes and suggests that Protenix’s confidence head needs improvement. Fig. S1C plots LDDT against the number of effective sequences (*N*^*eff*^, log-scale). We recover the classical trend—accuracy rises with *N*^*eff*^ —yet the slope is shallow.

Our benchmark confirms that protein monomer prediction has reached remarkable maturity across diverse folds and sequences, effectively solving what was once structural biology’s greatest challenge. Monomer prediction performance is relatively strong, as protein structure prediction has been extensively studied over the years. This advancement also provides a reliable basis for all protein-based interaction predictions in our study.

### Supplementary Note 2: Protein-peptide performance

**Fig. S2.**
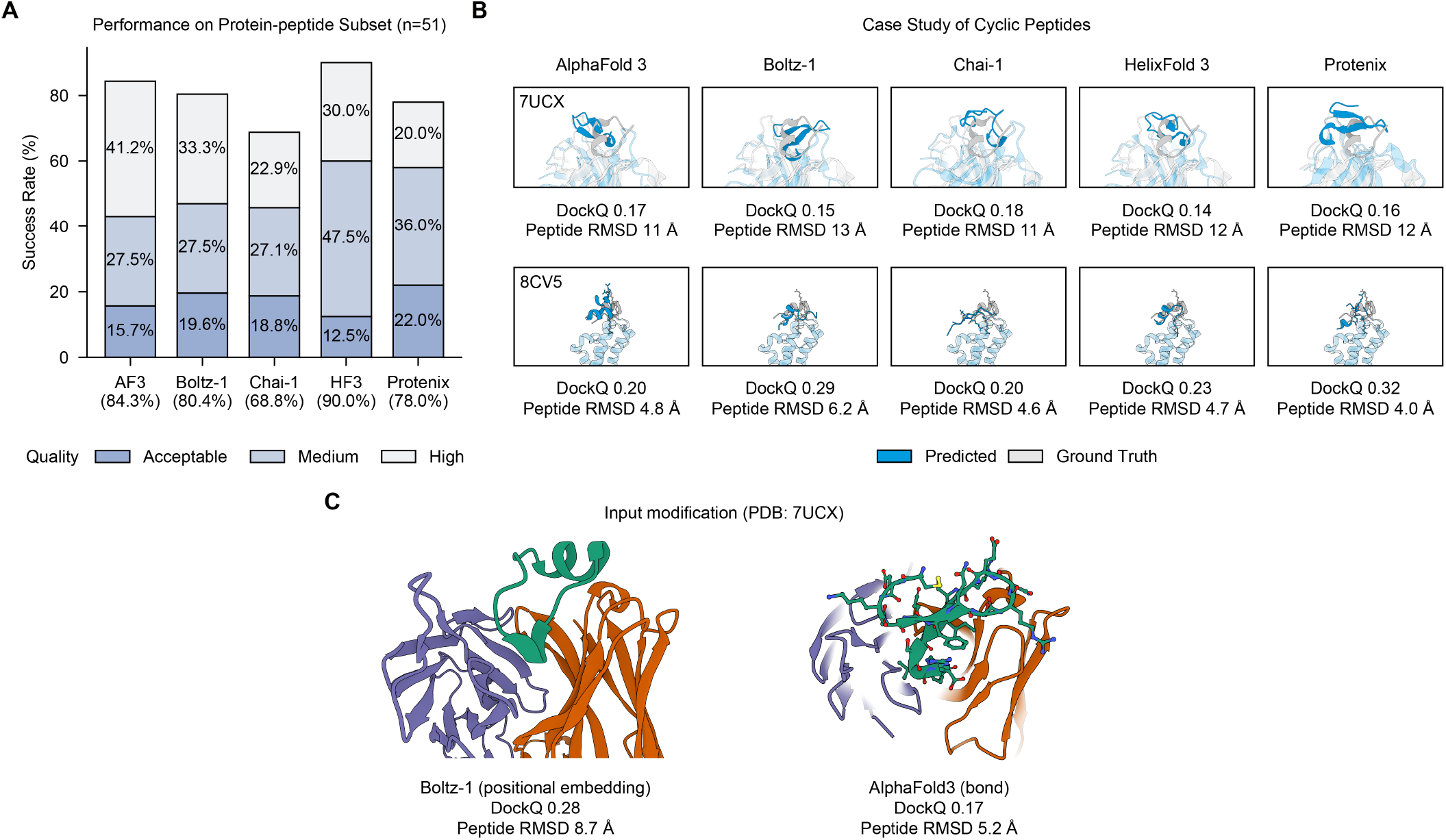
Model performance of protein-peptide interfaces. **A**. Docking performance on protein-peptide subset (n=51). **B**. Case study of two cyclic peptides. These cases are collected from the protein-protein subset since the peptide lengths are greater than 16 and classified as proteins in our benchmark. **C**. Case study of 7UCX with input modification.

Protein-peptide interactions mediate numerous signaling pathways and regulatory mechanisms through specific recognition of short linear motifs by specialized binding domains (1). These interfaces represent attractive therapeutic targets due to their compact binding footprints and potential for mimicry by small molecules. While current models performed well with structured peptides that bind to defined pockets, they struggle with flexible peptides that adopt specific conformations only upon binding. Accurate prediction of these interactions has significant implications for peptide drug design, targeting protein-protein interfaces, and understanding post-translational modification recognition, a critical aspect of cellular signaling networks.

Fig. S2A shows the performance on protein-DNA and protein-peptide interactions. Peptides are defined as amino acid chains with fewer than 16 residues. The dataset comprises 51 target interfaces. Overall, the models performed well, indicating that short peptides are relatively simple for the models to handle. HelixFold 3 was the best-performing model, achieving 90%, surpassing the other models. This might be attributed to proteins within this length range (<16 residues) representing the model’s “sweet spot”, a phenomenon consistent with observations from PepPCBench.(2)

Cyclic peptides represent a highly significant class of molecules in drug discovery due to their enhanced biophysical and pharmacological properties (3). Given the importance of cyclic peptides, we conducted an additional case study on a specific cyclic peptide structure (PDB: 7UCX, 8CV5). Fig. S2B displays the visualization results for the cyclic peptides 7UCX and 8CV5 from the benchmark set. For cyclic peptides, we identified two cases where the models did not perform well, showing a tendency to predict cyclic peptides as linear structures. The peptide RMSD values for the peptides were quite high, with the RMSD for 7UCX reaching around 11 Å. This indicates that there is still room for algorithmic optimization in cyclic peptide tasks. However, when the input is modified to provide more information about circular sequence of bonds (e.g. modifying positional encoding (4) for Boltz-1 v1.0.0 (https://github.com/jwohlwend/boltz/releases/tag/v1.0.0), and adding bond constraints for AlphaFold 3), the peptide can be predicted as a cyclic one and the structure prediction accuracy can be improved (Fig. S2C). The result suggests that improving the algorithms or acquiring more cyclic peptide data for training could enhance performance.

### Supplementary Note 3: Filtering of protein-ligand interfaces

Following the similar procedure established by PoseBusters (5), we applied quality control to the low-homology protein–ligand interfaces included in our benchmark. Ligand–protein pairs were retained only if they satisfied all of the following criteria:

1. The ligand molecular weight must lie between 100 Da and 900 Da.
2. The ligand must contain at least three heavy (non-hydrogen) atoms.
3. The ligand must only have H, C, O, N, P, S, F, and Cl atoms.
4. The parent structure must have a reported resolution better than 3.5 Å.
5. The ligand must not be covalently bonded to any other chain.
6. Polymer chains in the parent structure must not contain unresolved (unknown) residues.
7. The ligand real-space R-factor must be <0.2.
8. The ligand real-space correlation coefficient must be *≥* 0.95.
9. The ligand model completeness must be 100 %.
10. The PDB validation report must not list any stereochemical errors for the ligand.
11. The PDB validation report must not list any atomic clashes involving the ligand.
12. Initial ligand conformation could be generated using RDKit ETKDGv3.
13. The ligand SDF file must be parsable and sanitizable without errors using RDKit.
14. The minimum distance between the ligand and any protein atom must be at least 0.2 Å.
15. The minimum distance between the ligand and any other ligand in the complex must be at least 0.2 Å.
16. The minimum distance between the ligand and any metal ion in the complex must be at least 0.2 Å.
17. If a ligand has multiple protein interfaces within the same complex, only the interface with the smallest protein–ligand distance is retained.

**Table S1.** Counts of predictable and assessable targets on various tasks and models. Predictable refers to the count of generated CIFs, while Assessable indicates the quantity that can be scored by OpenStructure.

| Task | Subtask | Total | AlphaFold 3 |  | Boltz-1 |  | Chai-1 |  | Protenix |  | HelixFold 3 |  |
| --- | --- | --- | --- | --- | --- | --- | --- | --- | --- | --- | --- | --- |
|  |  |  | Predictable | Assessable | Predictable | Assessable | Predictable | Assessable | Predictable | Assessable | Predictable | Assessable |
| Protein Interaction | Ligand | 558 | 558 | 546 | 486 | 476 | 543 | 523 | 510 | 499 | 504 | 466 |
|  | Protein | 279 | 279 | 279 | 264 | 264 | 264 | 264 | 266 | 266 | 265 | 265 |
|  | Ab-Ag | 172 | 172 | 167 | 169 | 164 | 170 | 165 | 172 | 167 | 167 | 162 |
|  | DNA | 330 | 317 | 317 | 310 | 310 | 313 | 313 | 310 | 310 | 278 | 278 |
|  | RNA | 70 | 69 | 69 | 58 | 58 | 54 | 54 | 67 | 67 | 58 | 58 |
|  | Peptide | 51 | 51 | 47 | 51 | 48 | 48 | 45 | 50 | 48 | 38 | 36 |
| Monomer | Protein | 334 | 334 | 334 | 330 | 330 | 322 | 322 | 304 | 304 | 326 | 326 |
|  | DNA | 14 | 14 | 14 | 14 | 14 | 14 | 14 | 14 | 14 | 14 | 14 |
|  | RNA | 15 | 15 | 15 | 15 | 15 | 15 | 15 | 15 | 15 | 15 | 15 |

**Table S2.**
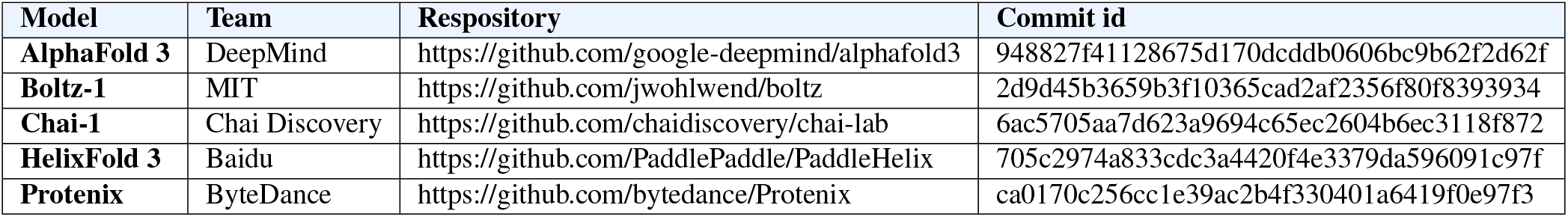
Details of different model inference pipelines.

**Table S3.**
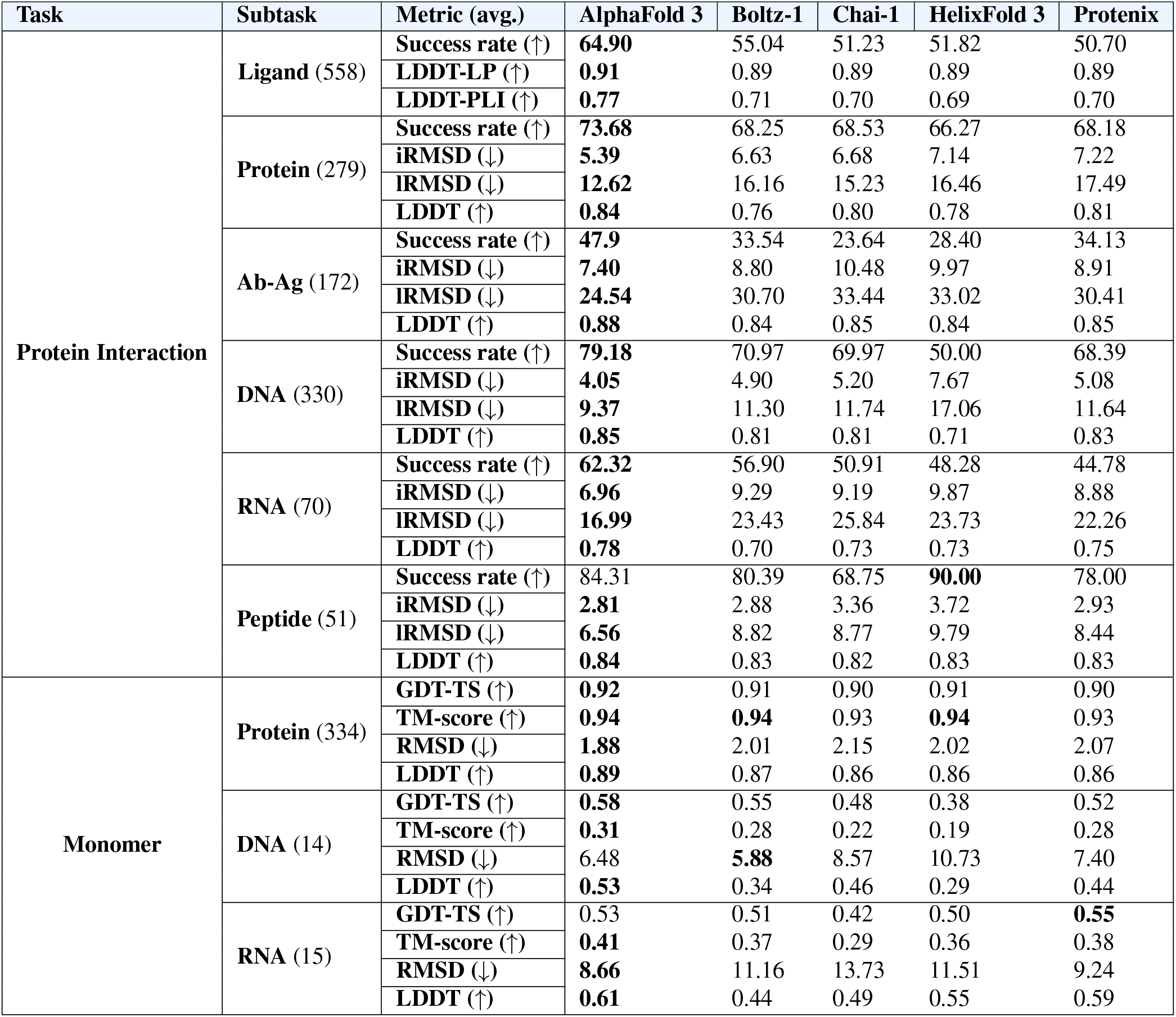
Detailed performance on each sub-task. Success rate is defined as the percentage of LRMSD < 2 Å and LDDT-PLI > 0.8 for protein-ligand interfaces and the percentage of DockQ score > 0.23 for other interfaces.

| Task | Subtask | Metric (avg.) | AlphaFold 3 | Boltz-1 | Chai-1 | HelixFold 3 | Protenix |
| --- | --- | --- | --- | --- | --- | --- | --- |
| Protein Interaction | Ligand (558) | Success rate (↑) | **64.90** | 55.04 | 51.23 | 51.82 | 50.70 |
| | | LDDT-LP (↑) | **0.91** | 0.89 | 0.89 | 0.89 | 0.89 |
| | | LDDT-PLI (↑) | **0.77** | 0.71 | 0.70 | 0.69 | 0.70 |
| | Protein (279) | Success rate (↑) | **73.68** | 68.25 | 68.53 | 66.27 | 68.18 |
| | | iRMSD (↓) | **5.39** | 6.63 | 6.68 | 7.14 | 7.22 |
| | | IRMSD (↓) | **12.62** | 16.16 | 15.23 | 16.46 | 17.49 |
| | | LDDT (↑) | **0.84** | 0.76 | 0.80 | 0.78 | 0.81 |
| | Ab-Ag (172) | Success rate (↑) | **47.9** | 33.54 | 23.64 | 28.40 | 34.13 |
| | | iRMSD (↓) | **7.40** | 8.80 | 10.48 | 9.97 | 8.91 |
| | | IRMSD (↓) | **24.54** | 30.70 | 33.44 | 33.02 | 30.41 |
| | | LDDT (↑) | **0.88** | 0.84 | 0.85 | 0.84 | 0.85 |
| | DNA (330) | Success rate (↑) | **79.18** | 70.97 | 69.97 | 50.00 | 68.39 |
| | | iRMSD (↓) | **4.05** | 4.90 | 5.20 | 7.67 | 5.08 |
| | | IRMSD (↓) | **9.37** | 11.30 | 11.74 | 17.06 | 11.64 |
| | | LDDT (↑) | **0.85** | 0.81 | 0.81 | 0.71 | 0.83 |
| | RNA (70) | Success rate (↑) | **62.32** | 56.90 | 50.91 | 48.28 | 44.78 |
| | | iRMSD (↓) | **6.96** | 9.29 | 9.19 | 9.87 | 8.88 |
| | | IRMSD (↓) | **16.99** | 23.43 | 25.84 | 23.73 | 22.26 |
| | | LDDT (↑) | **0.78** | 0.70 | 0.73 | 0.73 | 0.75 |
| | Peptide (51) | Success rate (↑) | 84.31 | 80.39 | 68.75 | **90.00** | 78.00 |
| | | iRMSD (↓) | **2.81** | 2.88 | 3.36 | 3.72 | 2.93 |
| | | IRMSD (↓) | **6.56** | 8.82 | 8.77 | 9.79 | 8.44 |
| | | LDDT (↑) | **0.84** | 0.83 | 0.82 | 0.83 | 0.83 |
| Monomer | Protein (334) | GDT-TS (↑) | **0.92** | 0.91 | 0.90 | 0.91 | 0.90 |
| | | TM-score (↑) | **0.94** | **0.94** | 0.93 | **0.94** | 0.93 |
| | | RMSD (↓) | **1.88** | 2.01 | 2.15 | 2.02 | 2.07 |
| | | LDDT (↑) | **0.89** | 0.87 | 0.86 | 0.86 | 0.86 |
| | DNA (14) | GDT-TS (↑) | **0.58** | 0.55 | 0.48 | 0.38 | 0.52 |
| | | TM-score (↑) | **0.31** | 0.28 | 0.22 | 0.19 | 0.28 |
| | | RMSD (↓) | 6.48 | **5.88** | 8.57 | 10.73 | 7.40 |
| | | LDDT (↑) | **0.53** | 0.34 | 0.46 | 0.29 | 0.44 |
| | RNA (15) | GDT-TS (↑) | 0.53 | 0.51 | 0.42 | 0.50 | **0.55** |
| | | TM-score (↑) | **0.41** | 0.37 | 0.29 | 0.36 | 0.38 |
| | | RMSD (↓) | **8.66** | 11.16 | 13.73 | 11.51 | 9.24 |
| | | LDDT (↑) | **0.61** | 0.44 | 0.49 | 0.55 | 0.59 |

**Fig. S3.**
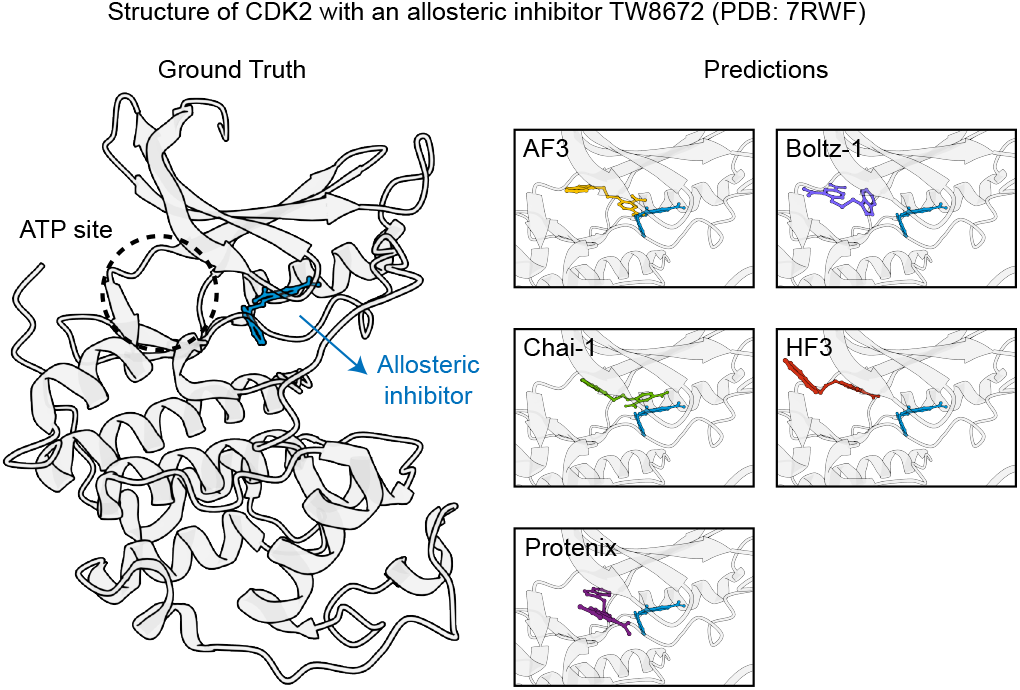
Case study of allosteric site prediction (PDB: 7RWF).

**Fig. S4.**
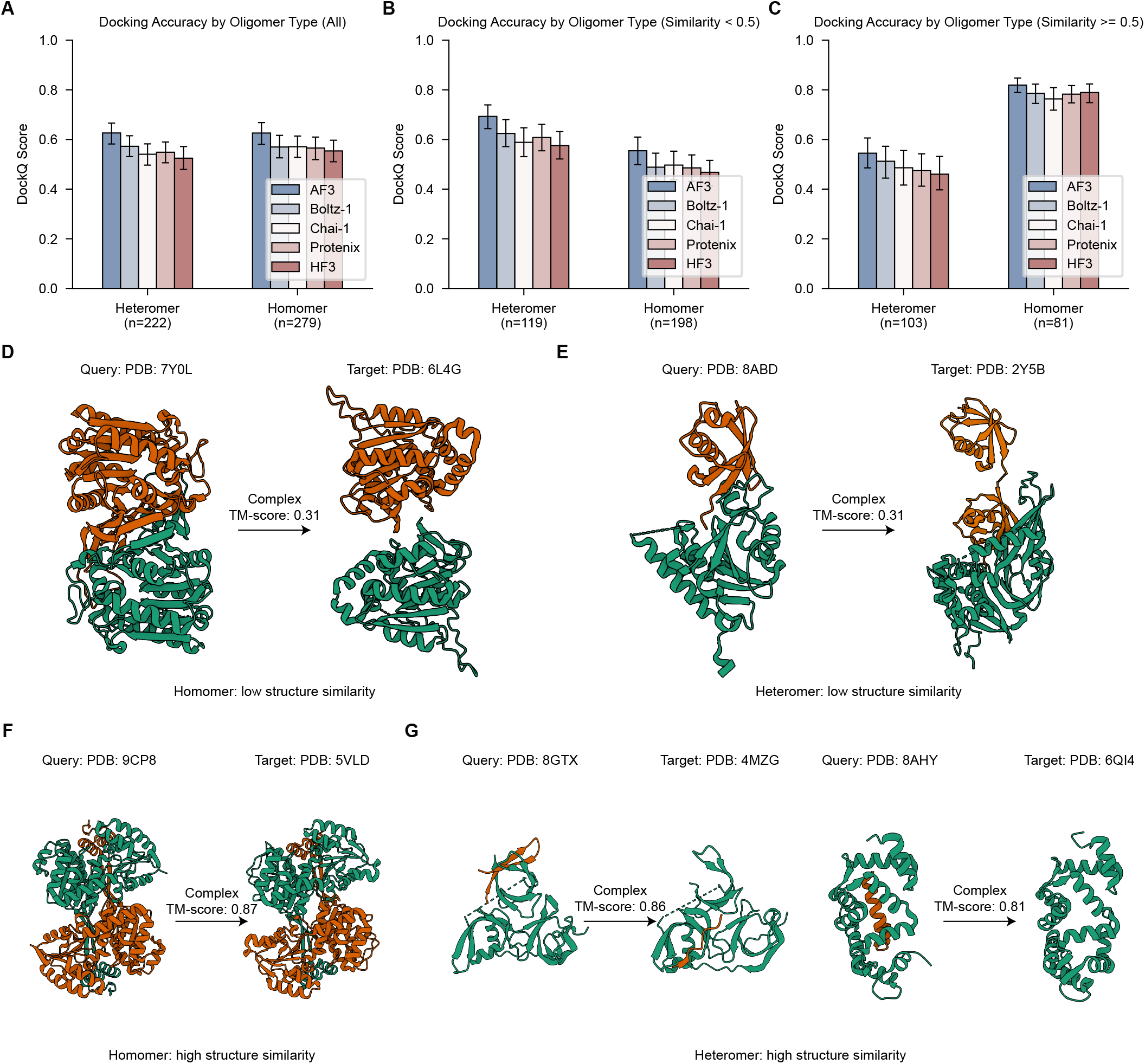
Protein-protein docking performance separated by oligomeric state. **A**. Targets at any similarity levels. **B**. Targets with structure similarity against the training set less than 0.5. **C**. Targets with structure similarity against the training set greater than or equal to 0.5. **D-E**. Representative low-similarity cases for homomer and heteromer. **F**. High-similarity homomeric pair. **G**. Two high-similarity heteromeric pairs whose global alignment fails to reproduce the correct interface, illustrating that high complex similarity does not guarantee successful docking orientation.

**Fig. S5.**
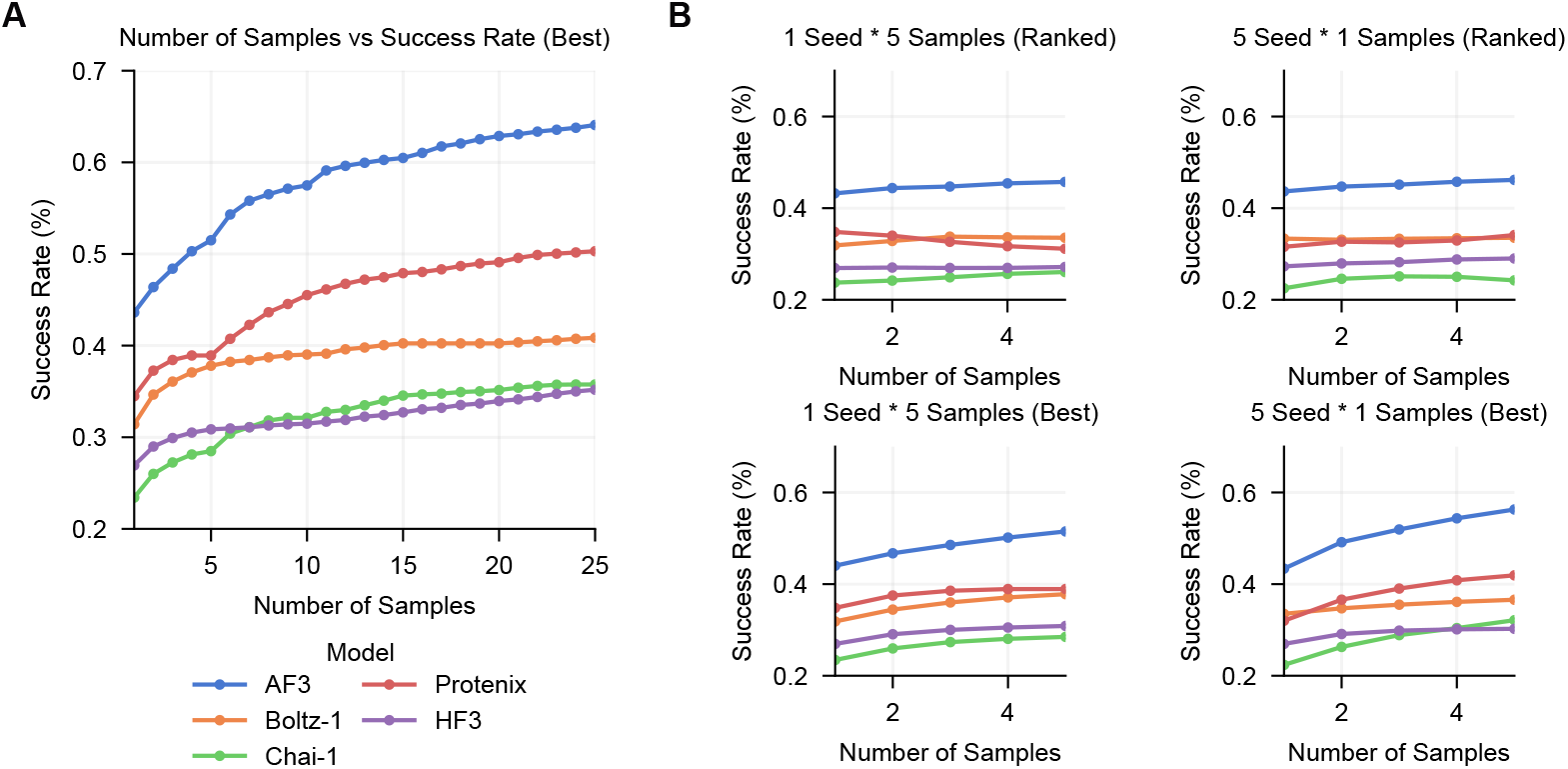
More correct antibody-antigen predictions can be found when the sample number increases. **A**. Selecting the top-scored sample from all the samples as the prediction. **B**. The individual impact of the number of seeds and samples on performance. The left side represents a fixed seed with an increasing number of samples, while the right side represents one sample per seed with an increasing number of seed predictions.

**Fig. S6.**
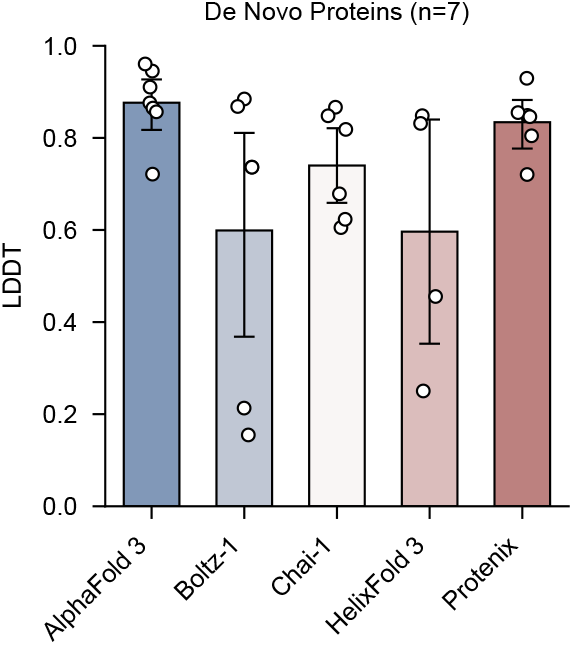
Performance on de novo protein monomers.

